# Effects of anti-RANKL, Zoledronate or combination therapy in a mouse model of fibrous dysplasia: a preclinical study

**DOI:** 10.1101/2025.09.08.672330

**Authors:** Biagio Palmisano, Chiara Tavanti, Giorgia Farinacci, Alessandro Corsi, Marta Serafini, Natasha M. Appelman-Dijkstra, Mara Riminucci

## Abstract

Bone fragility and pain are major clinical issues in fibrous dysplasia (FD) of bone, a genetic disorder characterized by increased bone resorption and lytic lesions. Both bisphosphonates (BPs) and denosumab are currently used to treat FD patients, although important concerns remain unsolved. BPs downregulate bone remodeling but their effects on FD lesions and pain are variable. Contrarily, denosumab converts FD tissue into mineralized bone and prevents disease progression, but disease rebound occurs upon treatment withdrawal. The combination of these two drugs may represent an effective and safe strategy for FD treatment.

We used a FD mouse model (EF1α-Gsα^R201C^) to assess whether zoledronate (ZOL) addition to anti-RANKL antibody (αRANKL) treatment could preserve the effects of RANKL inhibition after treatment discontinuation.

We show that αRANKL treatment rapidly reduced bone turnover markers (BTMs) and increased bone mass in affected skeletal segments, but FD lesions recurred shortly after discontinuation. Importantly, αRANKL+ZOL combination delayed disease rebound after αRANKL withdrawal, as bone density was preserved, BTMs rise was prevented, and no new lesions were observed. ZOL monotreatment increased bone density and reduced BTMs but did not fully halt disease progression. Finally, both αRANKL and αRANKL+ZOL treatments reduced fracture incidence and ameliorated pain-like behavior in FD mice.

These results demonstrate that combining zoledronate with denosumab may effectively treat FD. This strategy could particularly benefit patients with severe, rapidly progressive disease, in which RANKL inhibition would block lesion expansion and reduce bone turnover, while zoledronate would slow down the resumption of the disease and the rebound effect.

## Introduction

Inhibition of bone resorption is a widely used medical approach for patients with fibrous dysplasia of bone (FD, OMIM#174800), a skeletal disorder caused by activating mutations of the Gsα gene^1^. The rationale for the use of anti-resorptive medications in FD patients stems from the abnormally high bone remodeling at affected skeletal sites, where osteoclast formation and function are driven by Gsα mutated osteoprogenitor cells through RANKL secretion^2,3^. Indeed, uncontrolled bone resorption is one of the pathogenetic determinants that allow the development and expansion of the FD fibro-osseous tissue, which, in turn, severely subverts the architecture and mechanical properties of skeletal segments^1^. In FD patients, this pathological sequence can lead to bone deformity, fracture and pain. Ideally the anti-resorptive treatment in FD, as in other skeletal disorders with high bone remodeling, should not only halt osteoclast activity but also rescue the tissue pathology, thus providing long-term curative effect and likely, relief from pain. Furthermore, the treatment should not interfere with the skeletal growth in young patients neither it should cause undesired effects. Two classes of drugs are currently available that target different points of the molecular and cellular cascades linking osteoclast formation and activation to bone resorption, bisphosphonates (BPs) and denosumab. The former are structurally close analogs of pyrophosphate that bind to hydroxyapatite crystals and cause apoptosis of mature osteoclasts through different molecular mechanisms, depending on the absence or presence of nitrogen in their structure^4,5^. Denosumab is a humanized antibody that binds to RANKL favoring the apoptosis of pre-existing osteoclasts and inhibiting the generation of new ones^6^. In FD patients, both types of drugs are able to modulate bone turnover markers, improve bone mineral density and alleviate bone pain, although the analgesic function of BPs is not confirmed by all studies^7^. However, denosumab seems to have a higher ability to induce FD lesion refilling with bone and interfere with the progression of the disease^2,8,9^. This effect is supported by imaging assessment and bone biopsy^9,10^ in FD patients and multiple morphological analysis in FD mice, in which the replacing bone has been shown to be lamellar and hyper-mineralized^2,9,11,12^. The potential reversion of tissue pathology and the relatively reproducible effect on bone pain make denosumab the most promising medical treatment for FD. However, the disease rebound observed in some patients at treatment discontinuation^13^ favors a long-term administration that may have deleterious consequences on the skeleton, especially in pediatric patients. Sporadic studies in osteoporotic patients reporting that a timely administration of Zoledronate (ZOL), a potent third generation BP, may be efficacious in modulating bone biochemical markers after denosumab withdrawn^14,15^, suggest that a refined therapeutic schedule for FD based on both drugs could be a valid strategy to obtain a curative outcome while overcoming safety concerns. However, preclinical studies comparing the effects of RANKL inhibition, BPs, and combination therapy on FD lesions are still missing. Furthermore, it remains unclear whether or not BPs are able to preserve the histological changes induced by RANKL inhibition.

Using the EF1α-Gsα^R201C^ FD mouse model, we previously defined the minimum dose set of an anti-mouse RANKL monoclonal antibody (αRANKL) leading to radiographic and histological refilling of FD lesions, as well as the time interval required for the disease rebound after treatment discontinuation^2,8,12^. In this study, we asked whether this disease-free interval after αRANKL withdrawal could be extended by the addition of ZOL during the treatment. To address the point, the radiographic, biochemical and histological features of mice exposed to the combined treatment were compared to those of mice receiving either a continuous treatment with αRANKL or ZOL alone. Furthermore, using different behavioral tests previously validated on the same mouse model^16^, we also compared the effect of the different treatments on the mouse pain-like behavior.

## Methods

### FD mice and treatment

The study was conducted in compliance with relevant Italian laws and Institutional guidelines, and all procedures were IACUC approved. The FD mouse model was generated previously (EF1α-Gsα^R201C^)^17^. In this study, a total of 43 FD mice at 6 months of age were selected based on the presence of comparable radiographic skeletal phenotypes and randomly assigned to the different experimental groups. Both female (n=26) and male (n=17) mice were included in the study and, since no difference are known to occur in FD lesion appearance and development between the two sexes, data were pooled for the analyses. The anti-mouse RANKL monoclonal antibody (αRANKL, clone IK22/5, BioXcell, Lebanon, NH, USA) and Zoledronic acid (ZOL, Sigma Aldrich, Saint Louis, MO, USA) were injected intraperitoneally at 300 μg/mouse and 0.8 mg/kg, respectively. Both doses were formulated based on the conversion^18^ from the human doses of 60 mg for denosumab and 4 mg for ZOL considering a reference human body weight of 60 kg, i.e. the human doses were multiplied by 12.3. All mice were weighed at the beginning (T0) and the end (T12) of treatment.

Experimental groups are shown in Fig. 1. VEH group (n=8 mice) received the IgG control isotype (BioXcell) and PBS as vehicles. ZOL group (n=7 mice), received only two doses of ZOL. Mice in the other three groups received αRANKL as a loading dose (two injections) in the first week and then two additional doses, one in the second and one in the fourth week respectively, a dose set leading to conversion of FD tissue into bone in our model^12^. Then, the continuous αRANKL group (αRANKL-C, n=10 mice) received a dose of the antibody every three weeks until the end of the experiment; the discontinued αRANKL group (αRANKL-D, n=7 mice) did not receive any further doses and underwent a 9-week follow-up; the combination group (αRANKL+ZOL, n=11 mice) received two doses of ZOL, one during and the other after the αRANKL-D treatment.

**Figure 1.**
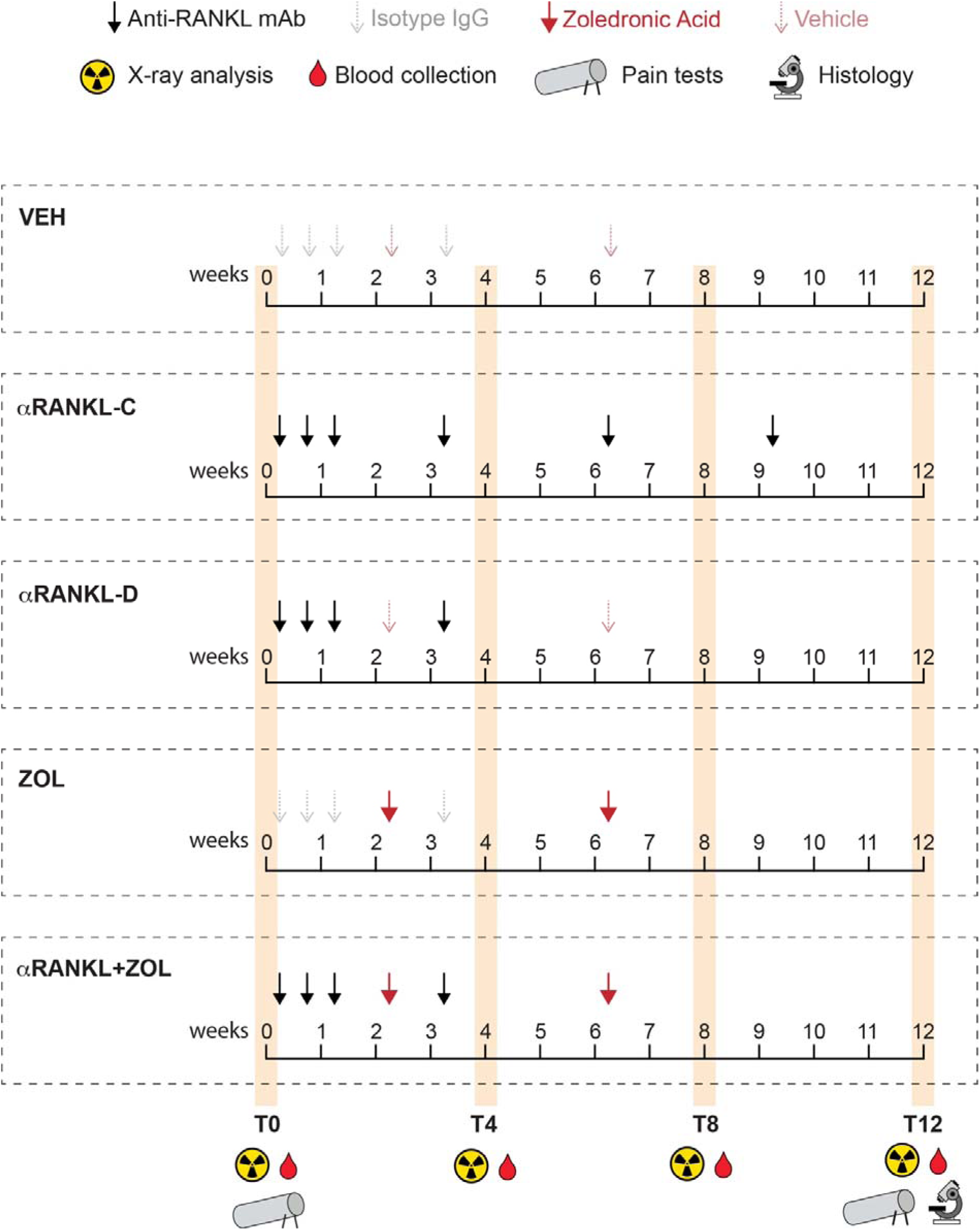
Scheme of the different treatment schedules.

All mice were sacrificed 12 weeks from the beginning of the experiments by carbon dioxide inhalation.

### Radiographic analysis and quantification of grey values

Whole body and tail radiographic analyses were performed under general anaesthesia before the beginning of treatment (T0) and after 4 (T4), 8 (T8) and 12 weeks (T12, Fig. 1), using Faxitron MX-20 Specimen Radiography System (Faxitron X-ray Corp., IL, USA) set at 25 kV for 8 sec and 24 kV for 6 sec for whole body and tail imaging respectively, with Kodak MINR2000 films. To evaluate changes in the radiodensity of bone segments, radiograms were scanned with an EPSON Perfection V750 PRO Scanner (Epson, Suwa, Japan) at 1200 dpi and saved as JPEG. Regions of interest (ROIs) were drawn in bone lesions of femurs or caudal vertebrae and radiodensity was evaluated by measuring mean grey values (MGVs) of each lesion^19^ using Adobe Photoshop 2024 (Adobe, CA, USA).

### Fracture count

One blinded observer counted fractures in the whole skeleton. We defined the fractures by evidence of solution of continuity and/or callus formation with or without bone deformity. In addition to the total number of fractures per group of treatment, we presented the number of new fractures that appeared during the experiment, visualized at each time point as the cumulative number developing during the treatment.

### Analysis of Bone Turnover Markers

Blood was collected from the mouse facial vein at the same time points at which radiographic analysis was performed (Fig. 1). Blood was allowed to clot for 30 min at room temperature and then centrifuged at 2000 ×g for 15 min at 4°C for serum separation. Sera were stored at -80°C until use. PINP and CTX-I levels were assayed using Rat/Mouse PINP EIA and RatLaps® (CTX-I) EIA (Immunodiagnostic system, UK) respectively, following manufacturer indications. Calcium and Phosphate levels were analyzed using QuantiChrom™ kits (BioAssay Systems, Hayward, CA, USA).

### Histology, histochemistry and histomorphometry

Mouse skeletal segments were collected after euthanasia at T12 (Fig. 1) and fixed with 4% formaldehyde at 4°C for 48 hours. Samples were then decalcified with 0.5 M EDTA at 4°C for 3–7 days and embedded in paraffin as per standard procedures. Three-μm-thick sections of femurs and lumbar vertebrae were used for histologic and histomorphometric analyses after hematoxylin/eosin (H/E) and sirius red/hematoxylin (SR/H) stains, and tartrate-resistant acid phosphatase (TRAP) histochemistry as previously reported^20,21^. Analyses were performed in a blinded fashion, using ImageJ software^22^ and following standard procedure and nomenclature^23^. The ROI was drawn 300 μm below the growth plate; H/E and SR/H-stained sections were used to measure the volume of trabecular bone per tissue volume (BV/TV), fibrous tissue per tissue volume (Fb.V/TV) and hematopoietic bone marrow per tissue volume (Ma.V/TV); TRAP-stained sections were used to measure osteoclast number per tissue area (N.Oc/TA) and mean osteoclast size (Oc. Size).

Quadriceps and spleens were dissected, weighted and fixed in 4% formaldehyde at 4°C for 48 hours. After paraffin embedding, transverse sections of the central region of muscles were cut and stained with H/E. The cross-sectional area (CSA) of muscle fibers was quantified using ImageJ^22^. Spleens were embedded in paraffin and longitudinal sections were obtained and stained with H/E and Perls’ Prussian blue (PPB) to detect iron deposits. Histomorphometry was performed on stained sections to assess the percentage of red pulp area over total tissue area (RP/TA) and the number of megakaryocytes per tissue area (N.MKs/TA) using ImageJ^22^.

### Behavioral tests

Burrowing and nesting tests were carried out as previously reported^16^ starting from 6 pm at specified time points (Fig. 1). A single investigator performed all procedures in a blinded manner, from animal allocation and handling to the endpoint behavioral measurements.

Burrowing test was carried out using a 20 cm long tube, with one end of the tube that was sealed with an aluminum lid and the other end that was left open and elevated about 3 cm off the floor. Habituation was carried in group by placing 4-5 mice into a 26 (length) × 23 (width) × 17 (height) cm cage containing the burrow tube filled with food pellets. Forty-eight hours later mice were individually transferred into a cage containing the tube filled with 300 grams of pellets for the beginning of the experiment. Burrowing activity was evaluated by weighing the remaining food in the tube after 2 hours (2H); at this time the tube was filled again with 300 grams of pellets and evaluation performed after an overnight experiment (ON).

For the nesting test, mice were transferred into individual cages containing 4.5 grams of shredded paper as nesting material without any other environmental enrichment. The next morning, the nest was photographed and given a score ranging from 1 to 5 based on the organization and the amount of paper used to build it. In cases where the results did not fit perfectly one of the scores a half point was either subtracted or added.

### Statistical analysis

A repeated measure (RM) one-way ANOVA with Dunnett’s multiple comparison test was used to detect statistical differences among the different time points of a specific treatment in MGV, fracture and BTM analyses. Ordinary one-way ANOVA with Dunnett’s multiple comparison test was used to compare the histological and behavioral endpoints among the different treatments. A two-way ANOVA mixed-effects model was used to detect statistical differences when both time and treatment were considered as independent variables, and a post-hoc Tukey’s test was used for multiple comparisons analysis. In behavioral experiments, a paired t test was used to compare the scores between T0 and T12 of each treatment. In all experiments a p-value value <0.05 was considered statistically significant and indicated with one or more asterisks for even smaller p-values. A p-value <0.1 (but greater than 0.05) was considered marginally significant and was reported in the graphs. Graphs and statistical analyses were performed using GraphPad Prism 10 (GraphPad Software, La Jolla, CA, USA).

## Results

### ZOL addition delays the radiographic recurrence of FD lesions after αRANKL treatment discontinuation

Radiographic monitoring performed at week 4, 8 and 12 on mouse femurs and caudal vertebrae showed that VEH mice developed new lesions during time and their disease progressed with tendency to skeletal deformities. In the VEH group, no overall differences were observed in the bone radiodensity of the femurs and caudal vertebrae (Fig. 2a-d). On the contrary, both αRANKL and ZOL, either alone or in combination, induced changes in regions affected by FD. However, the timeline and magnitude of changes were different depending on the drug and treatment schedule. In femurs, all RANKL-inhibited groups showed a rapid and homogeneous increase in the bone density of lesions, evident during the first radiographic assessment after 4 weeks of treatment (Fig. 2a, c). Thereafter, in the αRANKL-C group, which continued with one injection of αRANKL every three weeks, the induced changes in the bone segments were maintained throughout the experiment with no appearance of new lesions (Fig. 2a, c). In contrast, in the αRANKL-D group, in which treatment was stopped after 3 weeks, lytic lesions recurred, and the typical ground glass appearance was revealed by the imaging performed at week 8. (Fig. 2a, c). However, this was not observed if ZOL was added to this αRANKL schedule. In the combined treatment group (αRANKL+ZOL), the lesion opacity was preserved throughout the experiment and a general increase in radiodensity was also observed in the skeletal segments (Fig. 2a, c). Overall, the radiological effect of the combined treatment was comparable to that provided by the αRANKL-C schedule (Fig. 2c). In mice that received ZOL alone, radiographic changes were also observed albeit at a slower pace compared to the RANKL-inhibited mice. However, since they continued throughout the experiment, at week 12, the gain in radiodensity was comparable to the other treated groups, although small lytic areas persisted creating a non-homogeneous appearance (Fig. 2a, c).

**Figure 2.**
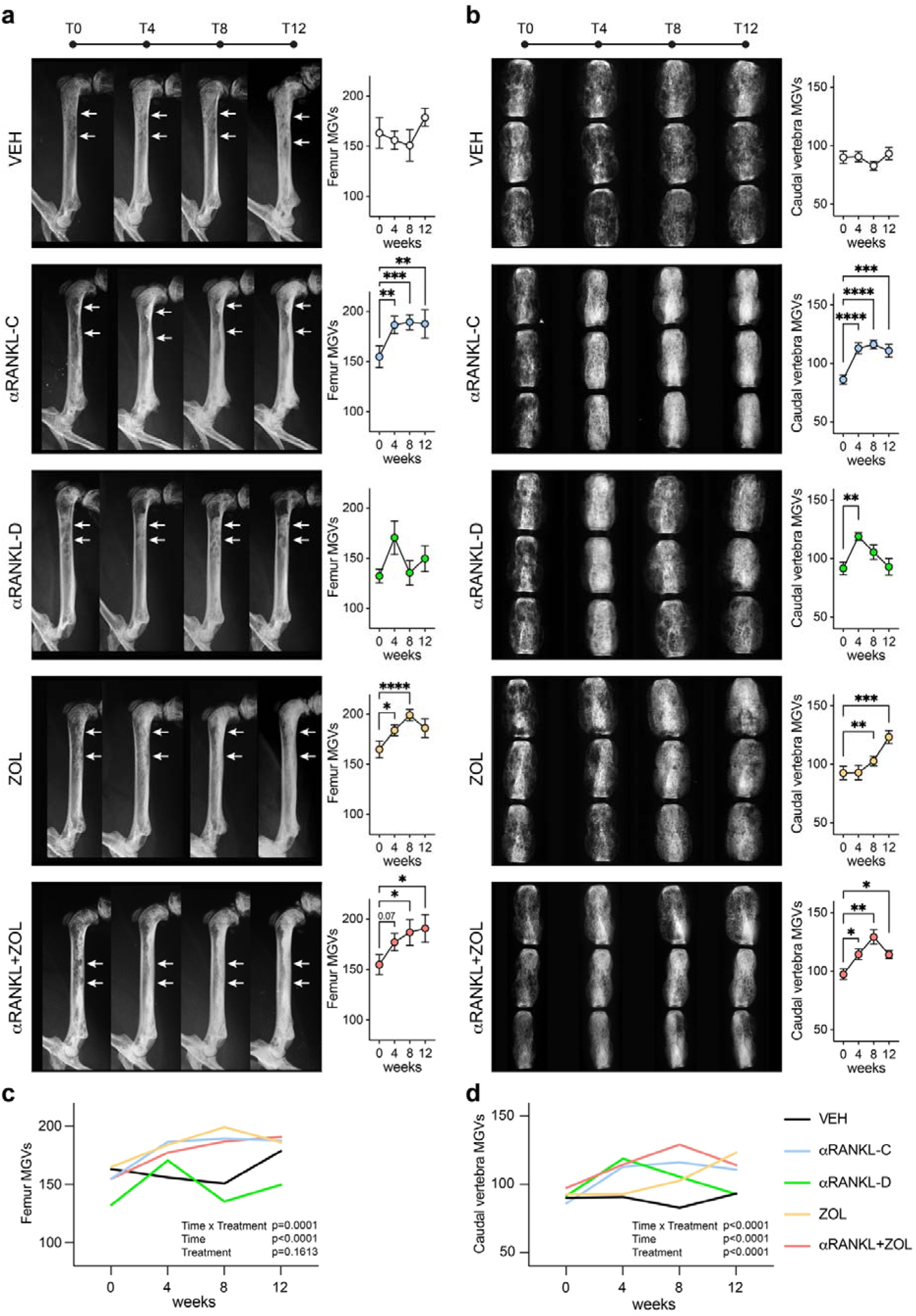
Radiographic effects of the different treatments on EF1α-Gsα^R201C^ mice. a, b) Radiographic analysis of the same femurs (a) and same caudal vertebrae (b) at different time points with radiodensity quantifications by mean grey values (MGVs). T4, T8 and T12 means were compared to T0 by RM one-way ANOVA Dunnett’s multiple comparison test. c, d) Grouped analysis of femoral (c) and vertebral (d) radiodensities during time in all experimental groups. P-values from the two-way ANOVA analysis are reported in the graph. In femur analyses, n=4-9 femur per time point. In caudal vertebra analyses n=15 vertebrae per time point. In all panels *p<0.05, **p<0.01, ***p<0.001, ****p<0.0001; p-value ≤0.1 is reported in the graphs.

In the tail vertebrae, the radiographic results were similar to those of the femurs (Fig. 2b). Again, an increase in lesional opacity was seen 4 weeks after RANKL inhibition was started and this was maintained throughout the active αRANKL treatment period. This treatment also prevented the abnormal enlargement of the tail vertebrae typically observed in untreated FD mice. As in the femurs, lesion recurrence with a ground glass appearance was detected in the αRANKL-D group at week 8 (Fig. 2b, d). The addition of ZOL maintained an overall higher bone density compared to αRANKL-D treated mice (Fig. 2d) and prevented the relapse of the disease after αRANKL discontinuation (Fig. 2b, d). In mice that received ZOL alone, vertebral radiodensity was markedly increased but the progression of the disease was revealed by ground glass changes and vertebral enlargement (Fig. 2b, d).

A two-way ANOVA was performed to analyze the effect of time and treatment on the MGVs. The analysis revealed a significant interaction between the effects of time and treatment in both femoral and vertebral MGVs (Fig. 2c, d). Simple main effects analysis showed that time but not treatment had a statistically significant effect on femur MGVs (Fig. 2c), whereas both time and treatment had a statistically significant effect on the density of caudal vertebrae (Fig. 2d).

### ZOL addition to αRANKL treatment reduced the incidence of fractures in FD mice

Radiographic analyses revealed that most FD mice showed at least one fracture at the beginning of treatment (Mean ± SD: 3.02 ± 2.22; median: 3). Most of the fractures were localized in the tail (92% of total fractures, Fig. 3a). We also observed fractures in long bones (7.2%) but rarely in the spine (0.8%). During the experiment, the VEH group had an increase in the number of new fractures, although comparisons among time points did not reach statistical significance (Fig. 3b). The groups of mice treated with αRANKL-D as well as ZOL alone showed a marked and statistically significant increase in the number of total fractures at the end of the experiment (Fig. 3b). In contrast, mice from the αRANKL-C and αRANKL+ZOL groups showed stabilization of the number of total fractures during the experiment (Fig. 3b). Accordingly, the analysis of new fractures developed after the beginning of treatment, revealed that while in αRANKL-D and ZOL groups an average of 2 new fractures was developed, in αRANKL-C and αRANKL+ZOL groups, the number of new fractures at week 12 was low and close to 0 (Fig. 3c). Two-way ANOVA analysis did not reveal a significant interaction between the effects of time and treatment on the count of new fractures (Fig. 3c). Simple main effects analysis showed that time but not treatment had a statistically significant effect on the appearance of the fractures (Fig. 3c).

**Figure 3.**
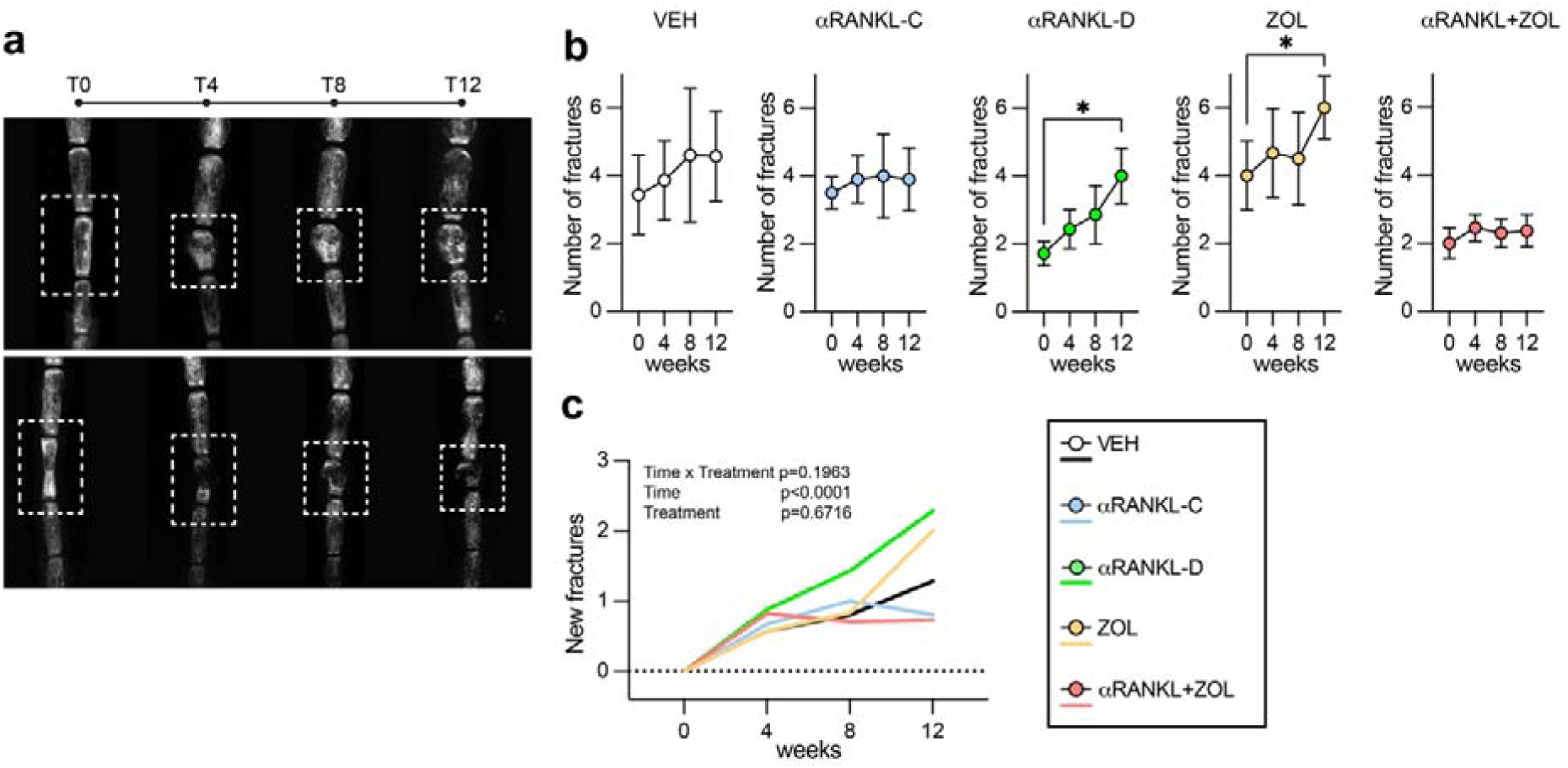
Fracture assessment in EF1α-Gsα^R201C^ mice following the different treatments. a) Representative radiographs of vertebral fractures in the tails of FD mice from αRANKL+ZOL (top panel) and VEH (bottom panel) groups. b) Count of the total number of fractures during time in each treatment regimen. One-way ANOVA with multiple comparison test, *p<0.05. T4, T8 and T12 means were compared to T0. c) Analysis of the number of new fractures after the beginning of treatment in each experimental group. The values at each time point represent the cumulative number of fractures. VEH n=8, ZOL n=7, αRANKL-C n=10, αRANKL-D, n=7, αRANKL+ZOL, n=11 mice. P-values from the two-way ANOVA analysis are reported in the graph.

### The addition of ZOL to αRANKL treatment prevents BTM rebound in mice after αRANKL discontinuation

To assess the effect of the different treatments on BTMs, we analyzed the serum levels of CTX-I and PINP, which reflect bone resorption and bone formation activity, respectively. In mice receiving VEH, CTX-I levels tended to increase during the treatment (Fig. 4a, b), while no changes in PINP levels were observed during time (Fig. 4c, d). In all mice receiving αRANKL, either alone or in combination with ZOL, CTX-I and PINP were markedly reduced at week 4 compared to baseline levels (Fig. 4a-d). In mice treated with αRANKL-C both CTX-I and PINP remained lower than baseline but did not decrease any further after 4 weeks (Fig. 4a-d). In the αRANKL-D group, BTMs levels started to rise after treatment cessation above baseline values (Fig. 4a-d). The observed rise was in parallel to the recurrence of lytic lesions and reached the values of VEH mice at week 12 (Fig. 4a-d). Interestingly, mice receiving the αRANKL+ZOL combined treatment, showed BTM changes similar to those of the αRANKL-C group, meaning a steep decrease in 4 weeks and plateau phase thereafter (Fig. 4a-d), and showed the lowest CTX-I level at all time points (Fig. 4b). In mice receiving ZOL alone, BTMs slowly declined during the treatment, resulting lower than VEH treated mice and comparable to the αRANKL-C and αRANKL+ZOL groups at week 12 (Fig. 4a-d). Two-way ANOVA analyses performed on CTX-I and PINP grouped data, confirmed the observed synergic effect of time and treatment, as the interaction between the two variables was statistically significant in both analyses (Fig. 4b, d). The Calcium levels did not show significant differences during time in each treatment group (Fig. 4e), while slight differences could be observed among the treatment groups (Fig. 4f), likely determined by different starting calcium levels. Phosphate levels were overall unchanged during time and among treatments (Fig. 4g, h), albeit an increase was noted at week 12 in all treatment groups (Fig. 4g, h).

**Figure 4.**
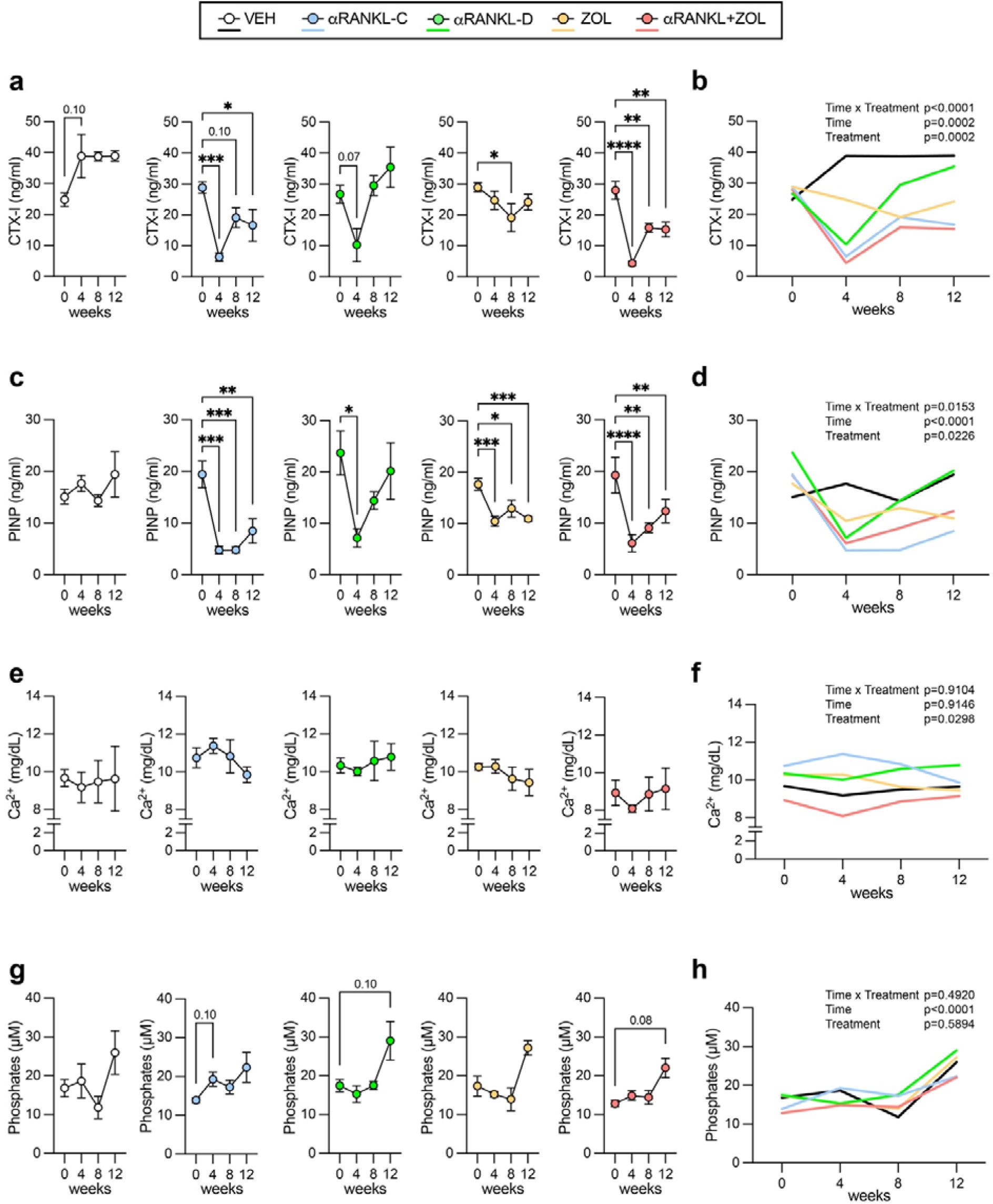
Bone turnover marker analysis at different time points in all treatment schedules. a) CTX-I levels in each treatment. b) Grouped analysis of CTX-I levels during time in all experimental groups. c) PINP levels in each treatment. d) Grouped analysis of PINP levels during time in all experimental groups. e) Calcium levels in each treatment. f) Grouped analysis of calcium levels during time in all experimental groups. g) Phosphates levels in each treatment. h) Grouped analysis of phosphates levels during time in all experimental groups. In a, c, e, g statistical analysis was performed with RM one-way ANOVA Dunnet’s multiple comparison test, and T4, T8 and T12 means were compared to T0. In b, d, f, h two-way ANOVA analysis was performed, and P-values are reported in the graph. In all the analyses, n=4-5 mice per time point. In all panels *p<0.05, **p<0.01, ***p<0.001, ****p<0.0001; p-value ≤0.1 is reported in the graphs.

### Combination of ZOL and αRANKL partially improved FD tissue pathology

All mice were sacrificed at T12 and histological and histomorphometric analyses were performed on femurs and lumbar vertebrae (Fig. 5 and S1). The VEH group showed all the pathological features of FD, with fibro-osseous lesions occupying the medullary cavity of affected segments (Fig. 5a and S1a). A similar percentage of bone and fibrous tissue was found in bone lesions of VEH mice by histomorphometry (Fig. 5b, c, f, g). Bone fraction was composed by a mixed woven and lamellar structure (Fig. 5a), and the fibrous tissue was accompanied by numerous osteoclasts (Fig. 5a, d, h).

**Figure 5.**
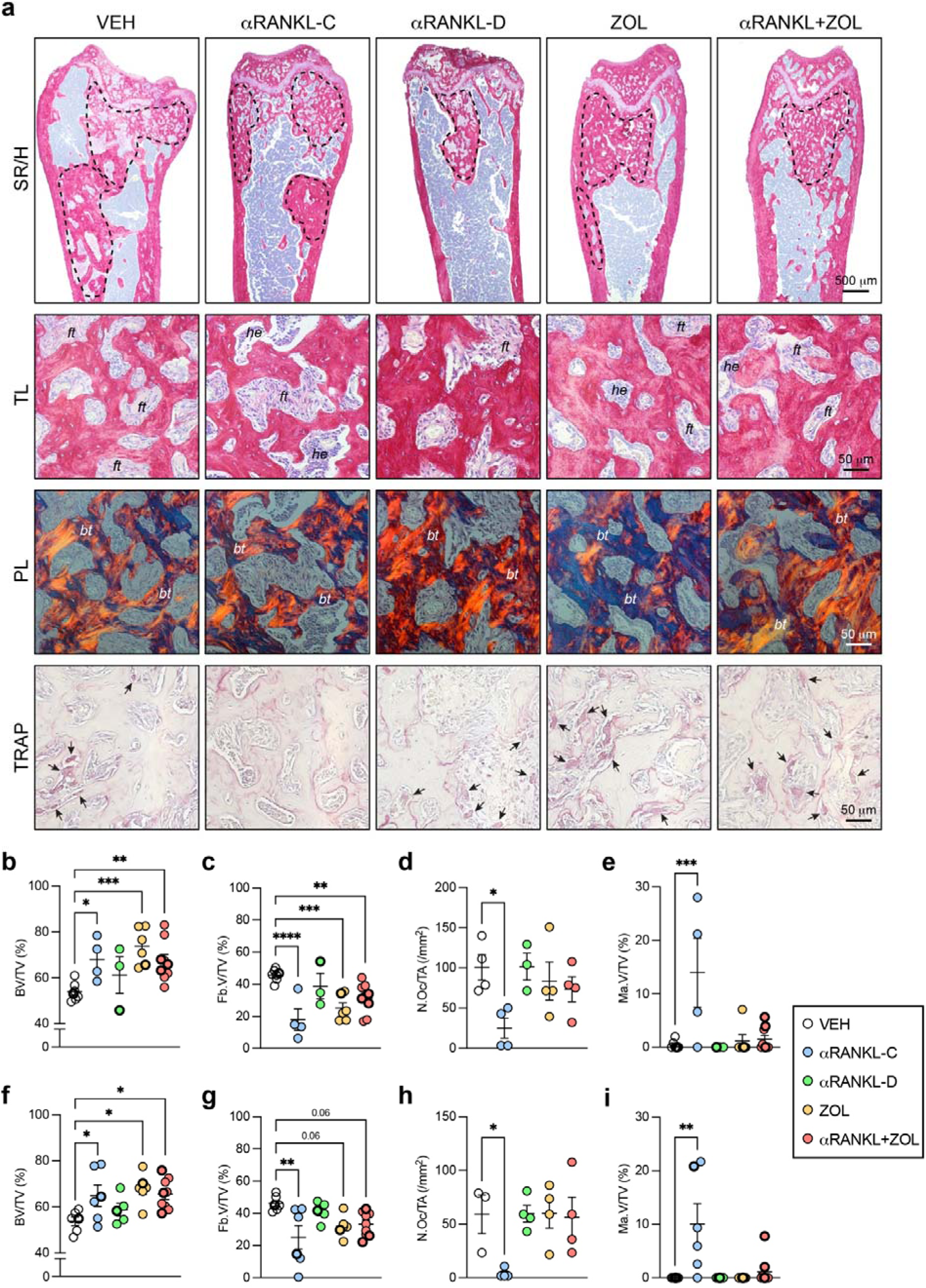
Histological and histomorphometric analyses of femurs and lumbar vertebrae in treated EF1α-Gsα^R201C^ mice. a) Representative sirius red/hematoxylin (SR/H)- and TRAP-stained sections of femurs with FD bone fibro-osseous lesions (dotted areas). Higher magnifications SR/H pictures are imaged with transmitted (TL) and polarized light (PL). *ft* = fibrous tissue; *he* = hematopoietic marrow; *bt* = bone trabeculae. TRAP pictures show red-stained osteoclasts (arrows). b-i) Histomorphometric analysis of intralesional bone volume (b, f), fibrous tissue volume (c, g), osteoclast number (d, h) and hematopoietic marrow volume (e, i) in femurs (b-e) and lumbar vertebrae (f-i). BV/TV: bone volume per tissue volume. Fb.V/TV: fibrous tissue volume per tissue volume. N.Oc/TA: number of osteoclasts per tissue area. Ma.V/TV: marrow volume per tissue volume. Data are presented as scatter plots showing all the experimental samples. Statistical analysis was performed using one-way ANOVA with Dunnett’s multiple comparison test where mean values from each experimental group is compared to VEH (control) group. In all graphs, each dot represents a data point. *p<0.05, **p<0.01, ***p<0.001, ****p<0.0001; p-value ≤0.1 is reported in the graphs. Differential labeling was used in the figure panels to distinguish data from female and male mice (i.e. males are represented with a thicker symbol border).

An increase in intralesional bone with a lamellar structure was observed in the αRANKL-C group (Fig. 5a, b, f and S1a). As expected, no differences compared to VEH were observed in the bone volume of mice in αRANKL-D group as these mice had already lost the bone mass that was previously achieved during the 3-week treatment course (Fig. 5a, b, f and S1a). A significant increase in bone mass compared to VEH was also observed in the group receiving the combination of αRANKL+ZOL and in the ZOL group (Fig. 5a, b, f, and S1b, a). The presence of fibrotic marrow was markedly reduced in the αRANKL-C group (Fig. 5a, c, g and S1a) as this was the only treatment able to significantly reduce the number of osteoclasts (Fig. 5a, d, h). Interestingly, histomorphometric analysis performed on areas previously occupied by FD lesions revealed that in this experimental group, an important fraction of marrow spaces among the newly formed bone was occupied by normal bone marrow filled with hematopoietic cells (Fig. 5a, e, i and S1a). In contrast, large lytic areas filled by fibrous tissue and abundant osteoclasts were detected in mice exposed to the αRANKL-D schedule comparable to VEH group and consistent with the radiographic recurrence of the disease after αRANKL discontinuation (Fig. 5a, c, g and S1a). Areas of fibrosis were also observed in both αRANKL+ZOL and ZOL groups but these were smaller compared to VEH (Fig. 5a, c, g and S1a). In αRANKL+ZOL and ZOL groups, numerous and giant osteoclasts were observed (Fig. 5a, d, h and S1b, c). As in the αRANKL-C group, a few mice receiving αRANKL+ZOL and ZOL showed small areas of hematopoietic tissue (Fig. 5a, e, i and S1a).

### RANKL inhibition is effective on mouse pain-like behavior

To evaluate the effect of the different treatments on mouse pain-like behavior, we performed the burrowing and nesting tests. Specifically, for each mouse we assessed the behavioral scores at baseline and immediately before the sacrifice (week 12, Fig. 1).

In the VEH group the burrowing and nesting behavior during the 12-week experiment declined (Fig. 6a, b, d, e), as test results worsened in most of the mice. In particular, the nesting behavior in VEH mice at T12 resulted significantly decreased compared to T0 by either paired t-test and two-way ANOVA (Fig. 6d, e). In mice receiving αRANKL, either alone or in combination with ZOL, a consistent amelioration of the pain-like behavior was observed (Fig. 6a-f). Burrowing behavior in αRANKL-D mice at T12 resulted significantly increased compared to T0 by paired t test (Fig. 6a), and delta change results of the same test revealed a significant increase in αRANKL-D compared to VEH (Fig. 6c). Similar results were observed in αRANKL-C group, although T0 vs T12 differences and delta changes results compared to VEH did not reach statistical significance (Fig. 6a-c). In the group that received αRANKL+ZOL combination therapy the burrowing and in particular the nesting capacity were considerably improved with delta changes resulting significantly increased than VEH (Fig. 6a-f). In ZOL treated mice, a worsening of mouse behavior was observed with time, and test scores were comparable to VEH suggesting that the ameliorating effect of the combination therapy was specifically due to αRANKL (Fig. 6a-f).

**Figure 6.**
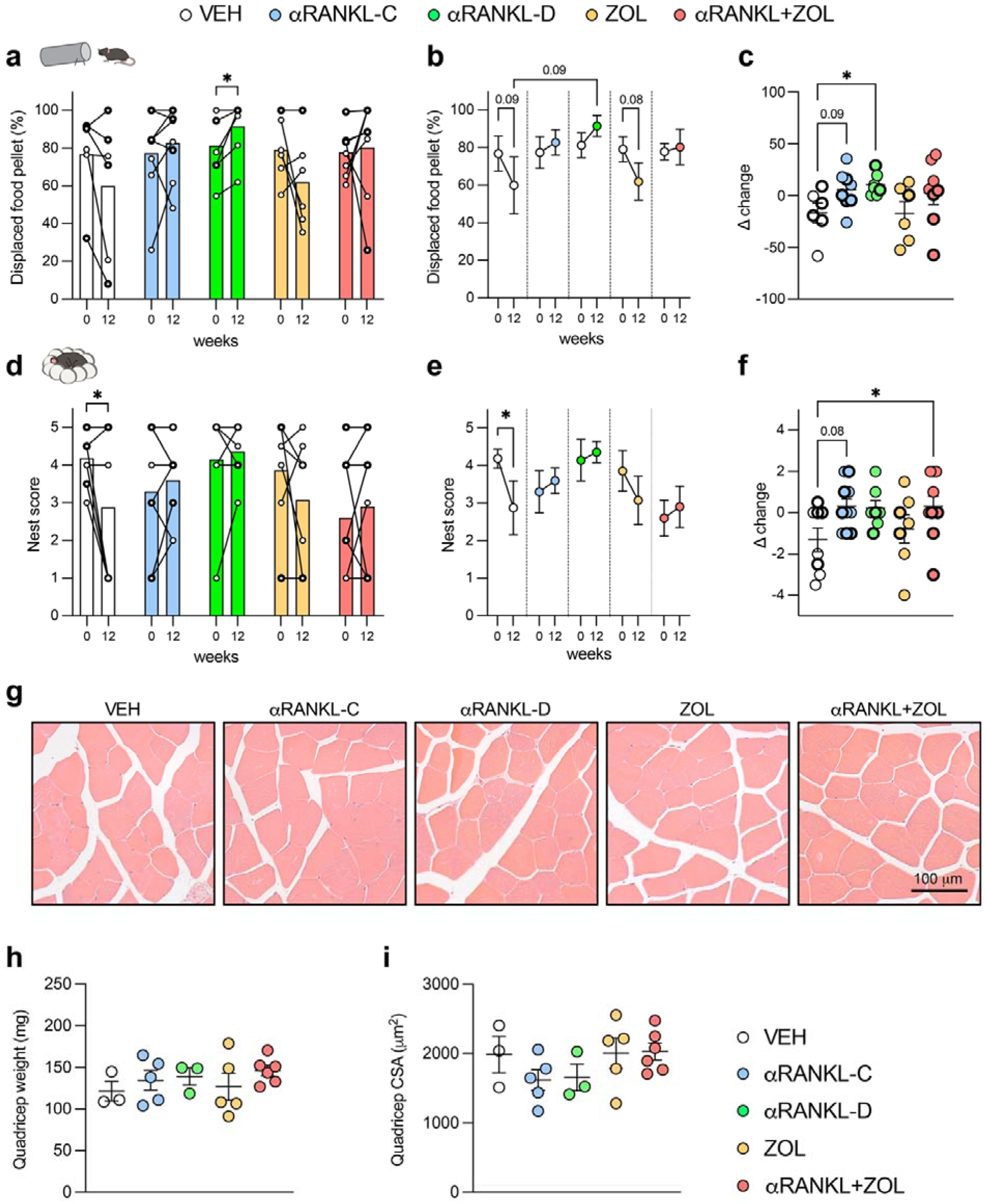
Assessment of pain-like behavior in treated EF1α-Gsα^R201C^ mice. a) Burrowing test results obtained before (T0) and at the end of treatment (T12) in the different experimental groups showing the behavior of each mouse during time. Statistical analysis performed with paired t test. b) The results reported in (a) are showed as mean ± SEM and analyzed with two-way ANOVA with Tukey’s multiple comparison test c) Delta change results calculated by subtracting T0 from T12 burrowing scores. Data above 0 indicates improvement in the behavioral result, while those below 0 reflect a behavioral worsening behavior during the treatment. Statistical analysis performed with one-way ANOVA. d) Nesting test results obtained before (T0) and at the end of treatment (T12) in the different experimental groups. Statistical analysis performed with paired t test. e) The results reported in (d) are shown as mean ± SEM and analyzed with two-way ANOVA with Tukey’s multiple comparison test. f) Delta change results calculated by subtracting T0 from T12 nesting scores. Statistical analysis performed with one-way ANOVA. *p<0.05; p-value ≤0.1 is reported in the graphs. g) Representative histological hematoxylin/eosin-stained sections of muscle fibers in mice at the end of treatments. h) Weight of quadriceps dissected from female mice at the end of treatments. i) Histomorphometric analysis of muscle fiber cross sectional area (CSA). In all graphs, each dot represents a data point. In a-f, differential labeling was used to distinguish data from female and male mice (i.e. males are represented with a thicker symbol border).

Of note, mice with severely reduced burrowing scores in basal conditions, i.e. burrowing less that 25% of the pellet food, did not show any amelioration, independent of the type of treatment (data not shown).

### Inhibition of bone resorption does not affect muscle fiber size

It has been recently suggested that RANKL inhibition may be beneficial for skeletal muscle function^24^. To understand if the improvement in the behavioral capacities seen in αRANKL-treated mice could be explained, at least in part, by improved muscular parameters, we performed a morphological evaluation of quadriceps (Fig. 6g). Muscle weight was similar amongst the different experimental groups at the end of treatment (Fig. 6h). Histomorphometric analysis of the CSA of the quadriceps revealed no differences among the groups, although a trend toward an increase of muscle fibers area was observed in αRANKL+ZOL-treated animals (Fig. 6i).

### ZOL administration induces splenomegaly and accumulation of megakaryocytes in spleens

We did not observe changes in the BW of mice in the different experimental groups, except for ZOL-treated mice, in which a significant increase was noticed (Fig. S2). At the time of the sacrifice, necroscopy showed that some mice had larger spleens (Fig. 7a). In particular, the spleens of mice receiving ZOL, either alone or in combination with αRANKL, were larger and heavier than those of VEH treated animals (Fig. 7a, b). Histological and histomorphometric analyses did not reveal differences in the iron deposits or amount of red pulp per tissue area among the different experimental groups (Fig. 7c, d), thus indicating the expansion of the entire splenic parenchyma rather than an unbalanced red/white pulp ratio. Interestingly however, an increase in the number of megakaryocytes often with clear intranuclear vacuoles was observed in the ZOL-treated animals compared to VEH group (Fig. 7e, f).

**Figure 7.**
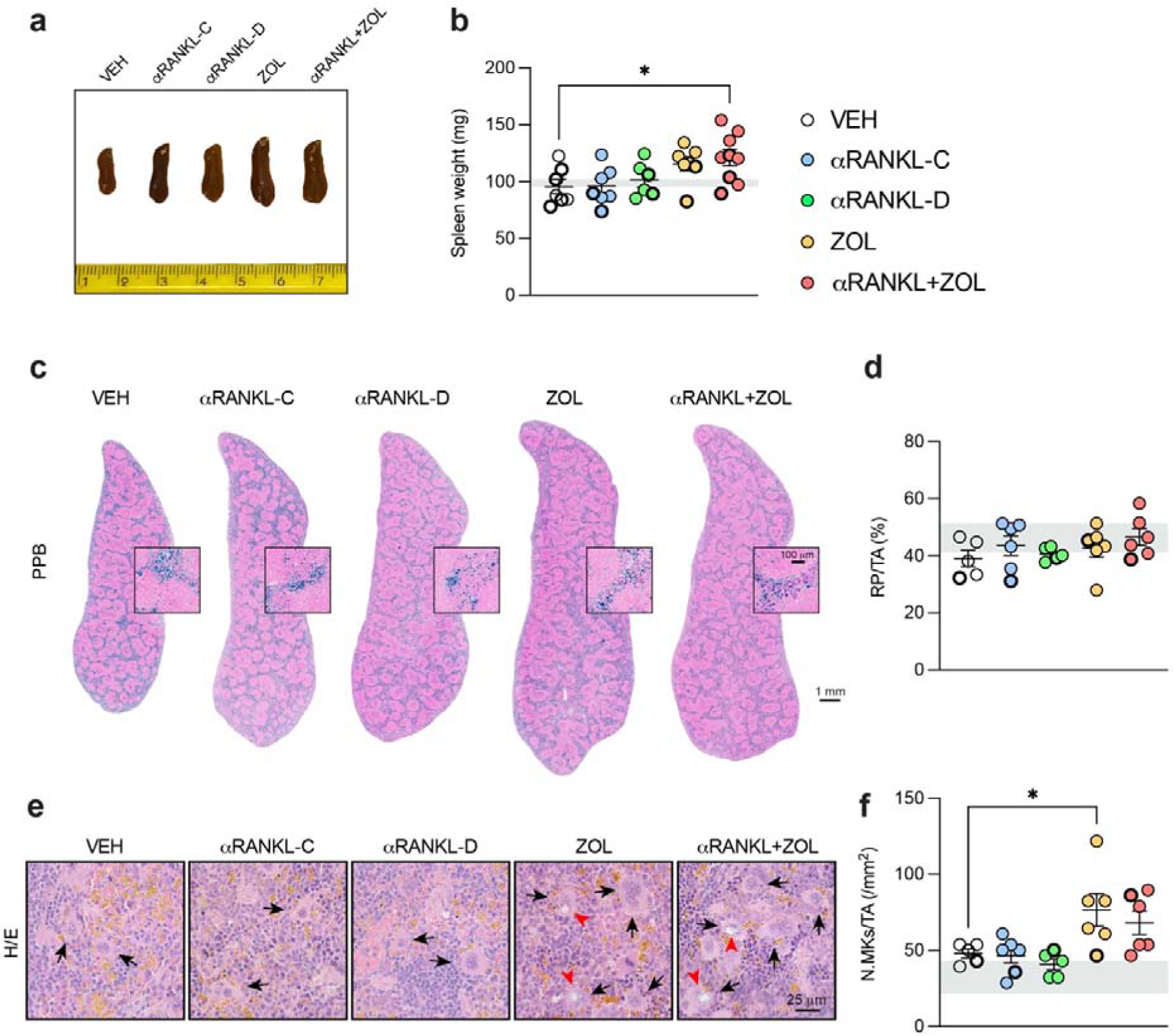
Assessment of spleen morphology in EF1α-Gsα^R201C^ mice at the end of the different treatments. a) Pictures of representative spleens from each group of treatment. b) Weight of spleen dissected from the mice at the end of treatments. c) Representative pictures of spleen sections stained with Perls’ Prussian blue (PPB) to detect iron deposits (blue-stained). d) Histomorphometric evaluation of red pulp over total tissue area (RP/TA). e) Picture of H/E-stained spleen sections showing megakaryocytes (arrows), some of them with intranuclear vacuoles (red arrowheads). f) Histomorphometric evaluation of megakaryocytes number over total tissue area (N.MKs/TA). Grey bands in the panels represent WT ranges for each specific parameter. In all graphs, each dot represents a data point. Differential labeling was used to distinguish data from female and male mice (i.e. males are represented with a thicker symbol border).

## Discussion

Discontinuation of denosumab raises a major clinical issue since post-treatment bone turnover activity is usually exaggeratedly increased compared to baseline and the bone mass gained during the treatment is rapidly lost^25^. In osteoporotic patients, this rebound effect associates with the risk of new vertebral fractures^26–29^ whereas in high bone remodeling diseases such as FD it may also be accompanied by severe hypercalcemia^13,30^. While theoretically feasible, long-term administration of denosumab is not recommended and strategies to preserve its effect after treatment cessation are highly needed. Clinical studies in osteoporosis suggest that administration of ZOL may be a reasonable approach, at least in patients who received a short-term denosumab treatment^14,15^ and The European Calcified Tissue Society (ECTS) recently proposed repeated administrations of this BP in patients with elevated BTM levels after denosumab discontinuation^31^. In order to start collecting proof-of-concept data that this strategy could be also effective in FD, we carried out a preclinical study on the EF1α-Gsα^R201C^ mouse model of the disease in which we compared different treatments including continuous αRANKL, discontinued αRANKL, ZOL, and αRANKL+ZOL combination. In the combined treatment, ZOL was administered during and after, rather than before, the αRANKL assuming that this would ensure a better incorporation into the newly formed, mineralized bone matrix than in the pre-existing FD bone, which is osteomalacic in both patients and mice^17,32,33^. Furthermore, the post-treatment dose was provided three weeks after the end of RANKL inhibition since, based on pilot studies, this is the optimal dosing interval to maintain the effect of αRANKL in long term treatments in EF1α-Gsα^R201C^ mice.

We show that ZOL was effective in preserving the radiographic improvement of femoral FD lesions for at least 9 weeks after the last dose of αRANKL, with an overall level of bone density that was comparable to that of mice in the αRANKL-C group. Of note, a significant increase in the radiodensity of affected segments was also observed in mice receiving ZOL alone, in agreement with previous studies reporting radiological amelioration in pediatric FD patients treated with this BP^34^. However, compared to both the αRANKL-C and the combined groups, bone mass accrual in ZOL receiving mice proceeded slower and led to more marked bone enlargement/deformities. This can be reasonably explained by the different mechanism of action of the two drugs. By completely blocking bone resorption, αRANKL allowed the rapid conversion of the fibrous tissue into bone^2,8,9^, after which pathological bone was no longer formed. In contrast, during ZOL treatment, pathological bone continued to be deposited while its resorption was reduced due to the downregulation of osteoclast function. Furthermore, despite an increase in radiodensity, ZOL alone did not prevent the occurrence of new fractures, mainly occurring in the caudal vertebrae of these mice. This observation is in agreement with the clinical evidence that in FD, radiographic amelioration induced by BPs is not predictive of a lower fracture risk^35^ as also observed in osteoporosis^36^. In contrast, mice that received the αRANKL+ZOL schedule showed the lowest incidence of fractures, as those in the αRANKL-C group, suggesting that the addition of ZOL may also preserve the beneficial effect of RANKL inhibition on this clinical complication^37^.

At the biochemical level, ZOL prevented the rise of BTMs observed after αRANKL discontinuation, maintaining their levels comparable to those of αRANKL-C mice at all time points. Consistent with reports on FD patients^38^, a reduction of BTMs was also observed in mice receiving ZOL alone, especially at weeks 8 and 12. However, the difference between the pre- and post-treatment level was smaller than that observed in both the αRANKL-C and the combined groups. Altogether, these data confirm that in FD, αRANKL administration provides a rapid improvement of radiographic features and BTMs and suggest that ZOL may be a good option to maintain these results over time. Since mice were monitored for 9 weeks after the last dose of αRANKL, it can be predicted that in FD patients receiving a similar schedule the effect of ZOL should last more than 5 years^39^. However, further studies are required to assess how this schedule can be translated into clinical settings.

We previously showed that a rise in calcium levels in young EF1α-Gsα^R210C^ mice occurred after 14-week treatment with high cumulative dose of αRANKL^2^. In this study, in which the total amount of αRANKL and the treatment duration were much lower than in the previous study, we did not observe any case of hypercalcemia. This suggests that hypercalcemia after discontinuation of αRANKL may occur in long duration, and therefore high cumulative dose, treatments, a hypothesis that seems to be corroborated by recent clinical studies^30,40^. Furthermore, since our previous study used very young mice^2^ whereas the current study treated adult mice, the observed hypercalcemia may be age-dependent—consistent with rebound hypercalcemia being more common in young patients following denosumab cessation^30,41^.

Consistent with the radiographic results, histomorphometric analyses performed at the end of the experiment showed a higher bone mass compared to controls in all the experimental groups, except for mice that underwent a short RANKL inhibition. As previously reported, continuous administration of αRANKL was the only treatment able to reduce significantly the amount of fibrous tissue and restore the hematopoietic microenvironment^2^. However, a decrease in the volume of fibrous tissue was also observed in mice receiving ZOL alone. The notion that ZOL in FD can partially reduce the fibrous tissue was never reported before and may be taken as further evidence that osteoclast activity plays a major role in the tissue pathology of FD. Typical histological feature of ZOL treatment, either with αRANKL or alone, was the presence of giant osteoclasts, as already noticed in the same transgenic mouse model of FD^8^. Giant osteoclasts following treatment with nitrogen containing BPs were initially described in postmenopausal women as cells with extended lifespan and apoptosis and poor bone resorbing capacity^42^. Although we did not investigate the mechanism of formation of giant osteoclasts, the strong decrease in the bone resorption marker CTX-I in the αRANKL+ZOL and ZOL groups confirmed that even in FD giant osteoclasts have a reduced bone resorbing activity and this may contribute to the positive effect on intra-lesional bone mass exerted by this BP.

To add more clinical perspective to our work we also performed specific tests to assess mouse pain-like behavior at both the beginning and end of each treatment. We observed that the test scores were ameliorated by αRANKL, either alone or in combination with ZOL. Pain improvement by denosumab was reported in many FD patients^13,43,44^ and it was speculated to be based on the reduction of the acidic environment caused by complete ablation of osteoclasts. This hypothesis is in agreement with our work, as both continuous administration of αRANKL and combination therapy with ZOL induce ameliorating effects on the pain-like behavior of treated mice. However, we also show that behavioral scores were improved in αRANKL-D mice, in which rebound had occurred, osteoclasts had regenerated, and FD had already relapsed. This result could be hardly reconciled with a role for the acidic environment in FD pain. Nonetheless, it is plausible that a short RANKL inhibition may still be sufficient to generate an early analgesic effect, as reported in patients with low back pain^45^, and a longstanding response. Of note, in mice with severe behavioral impairment at baseline, no amelioration was observed at the end of the experiment, independent of specific αRANKL treatment. This finding may explain the absence of significant analgesic effects of denosumab in some FD patients. More important, it suggests that the treatment should be administered soon after the appearance of pain in order to obtain a positive response. As in clinical settings, ZOL did not affect the behavioral score as a behavioral deterioration comparable to that of VEH mice was observed in the time frame of the experiment. Since recent studies demonstrated that αRANKL can produce beneficial effects on muscle mass and function in mice^24^ (although this is still a controversial issue in humans^46^), we also analyzed skeletal muscle morphology. We did not observe significant differences in quadricep weight, nor in the CSA of muscle fibers among the different groups. A slight tendency to larger muscle fibers was observed in ZOL treated mice, a result already reported in other mouse models of high bone remodelling^47^. However, ZOL did not ameliorate pain-related behavior, thus ruling out a potential association between muscle structure and function and the results of the behavioral analyses.

Finally, we found that mice receiving ZOL, either in combination with αRANKL or alone, had larger spleens with morphologically abnormal megakaryocytes compared to the other experimental groups. It was previously reported that in mice injected with a nitrogen-containing BP (namely the AHBuBP) the spleen was considerably larger than in mice receiving a chloro-containing BP and controls^48^. BP-dependent splenomegaly can be ascribed to the expansion or the red pulp with an increased number of megakaryocytes as well as to the development of extramedullary erythropoiesis in parallel with the depletion of bone marrow-resident macrophages^48^. In addition, it is known that nitrogen-containing BPs accumulate for several days into liver and spleen^49^ after the formation of complexes with calcium and iron^5^. We observed an increase in megakaryocytes but no difference in the amount of iron in the spleen in our ZOL-treated mice compared to the other experimental groups, thus we did not identify a potential pathogenetic mechanism. Further investigation is required to clarify this point. However, it is important to remark that to date, splenomegaly upon treatment with nitrogen containing BPs has been observed only in mice and therefore we cannot exclude that this side effect may be species-specific, resulting, for example, from the presence of physiological hematopoietic activity in the mouse but not human spleen.

In conclusion, we report that in a mouse model of FD, ZOL addition during and after RANKL inhibition was able to preserve the intra-lesional bone mass gain and to maintain the effect on BTMs and fracture risk for at least 9 weeks after αRANKL discontinuation. We also show that ZOL alone was able to increase bone mass but did not prevent the expansion/deformity of affected skeletal segments neither reduced the incidence of bone fractures and pain. Translated into the clinical scenario, the combination of denosumab with ZOL may be a useful approach especially in FD patients with a very active and rapidly progressive FD in which a short course of RANKL inhibition may be necessary to block lesion expansion/bone deformities and to relieve bone pain whereas ZA may be useful to slow down the resumption of the disease.

## Acknowledgments

This study was supported by grants from the Orphan Disease Center University of Pennsylvania in partnership with Fibrous Dysplasia Foundation (MDBR-19-110-FD, MDBR-21-110-FD, MDBR-23-010-FDMAS) to M.R.; the European Calcified Tissue Society (ECTS, Basic/Translational Research Fellowship 2021) to B.P.; Sapienza University to B.P. (AR223188B3DFA88B), and to A.C. (RP124190960A2E07).

The authors are grateful to the European Association Friends of McCune-Albright Syndrome (EAMAS) for the support (donation).

## Contributions

B.P. and M.R. designed the research study and drafted the manuscript. B.P. conducted the experiments, acquired and analyzed the data, and oversaw all aspects of the study. C.T. and G.F. conducted the experiments and acquired histological data. A.C., M.S. and N.A.D. provided expertise related to the experiments and contributed to the interpretation of the results. B.P., A.C., M.S., N.A.D. and M.R. edited and revised the manuscript. All the authors read and approved the manuscript.

---

**Figure S1.**
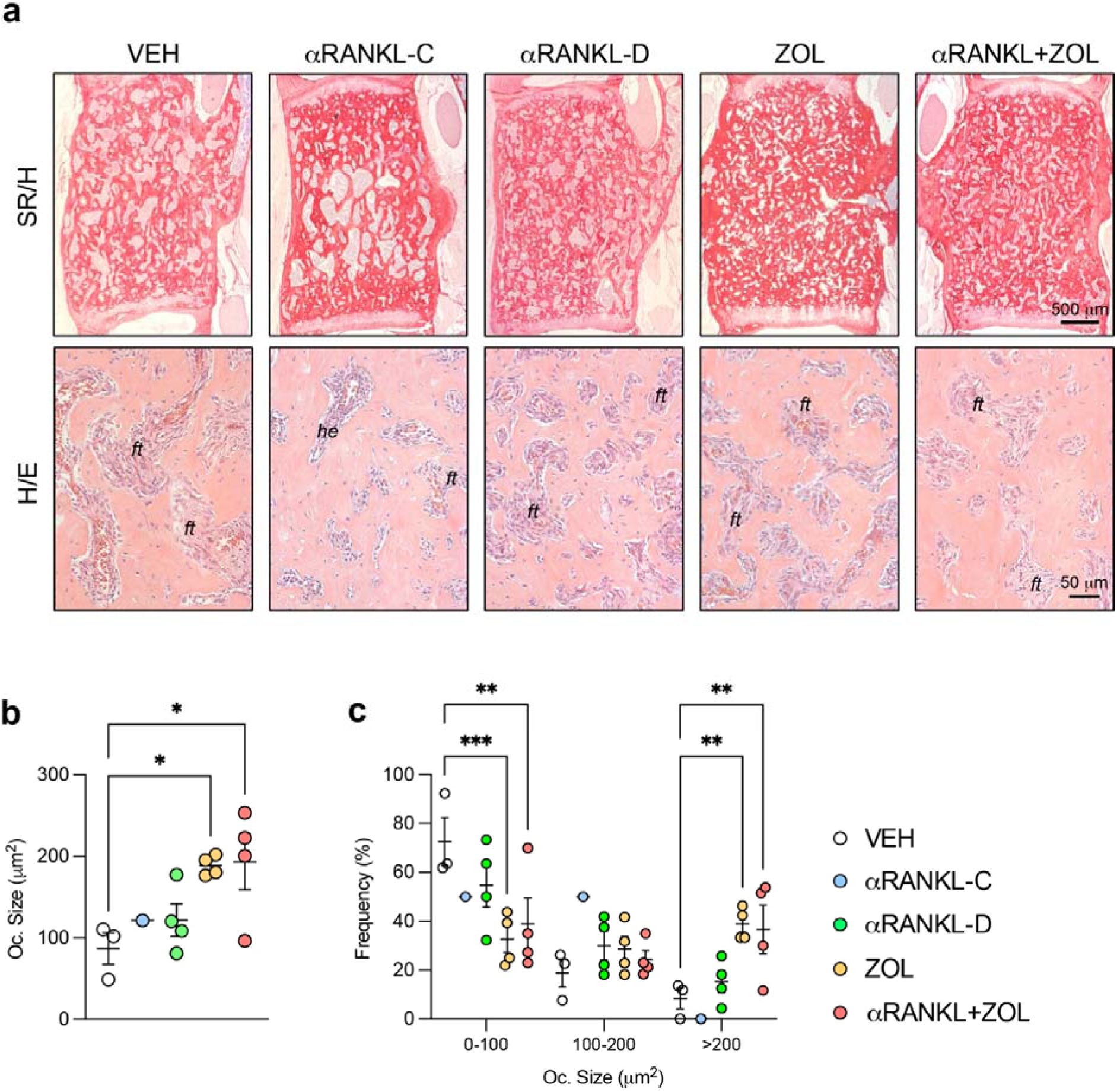
Histological and histomorphometric analyses of lumbar vertebrae in treated EF1α-Gsα^R201C^ mice. a) Representative SR/H- and H/E-stained sections of lumbar vertebrae with fibro-osseous lesions. *ft* = fibrous tissue; *he* = hematopoietic marrow. b) Histomorphometric evaluation of osteoclast size (Oc. Size) in female mice. Statistical analysis with one-way ANOVA and Dunnett’s multiple comparison test. c) Frequency of distribution of different osteoclast size range. Two-way ANOVA and multiple comparison test. *p<0.05, **p<0.01, ***p<0.001.

**Figure S2.**
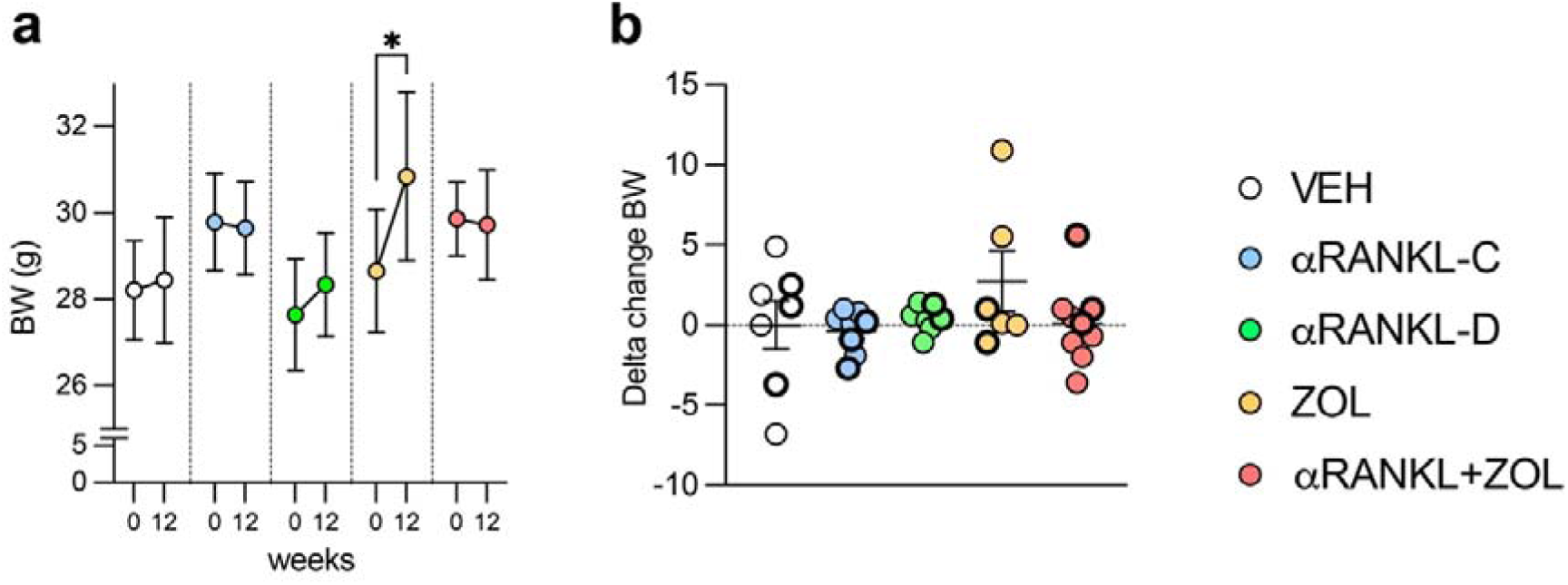
Body weight changes during the treatment in EF1α-Gsα^R201C^ mice. a) Body weight values at the beginning (0) and the end (12) of treatment. Two-way ANOVA with Tukey’s multiple comparison test. b) Delta change results calculated by subtracting T0 from T12 body weight.

