## Supplementary figures for "Effects of anti-RANKL, Zoledronate or combination therapy in a mouse model of fibrous dysplasia: a preclinical study"

**Figure S1**

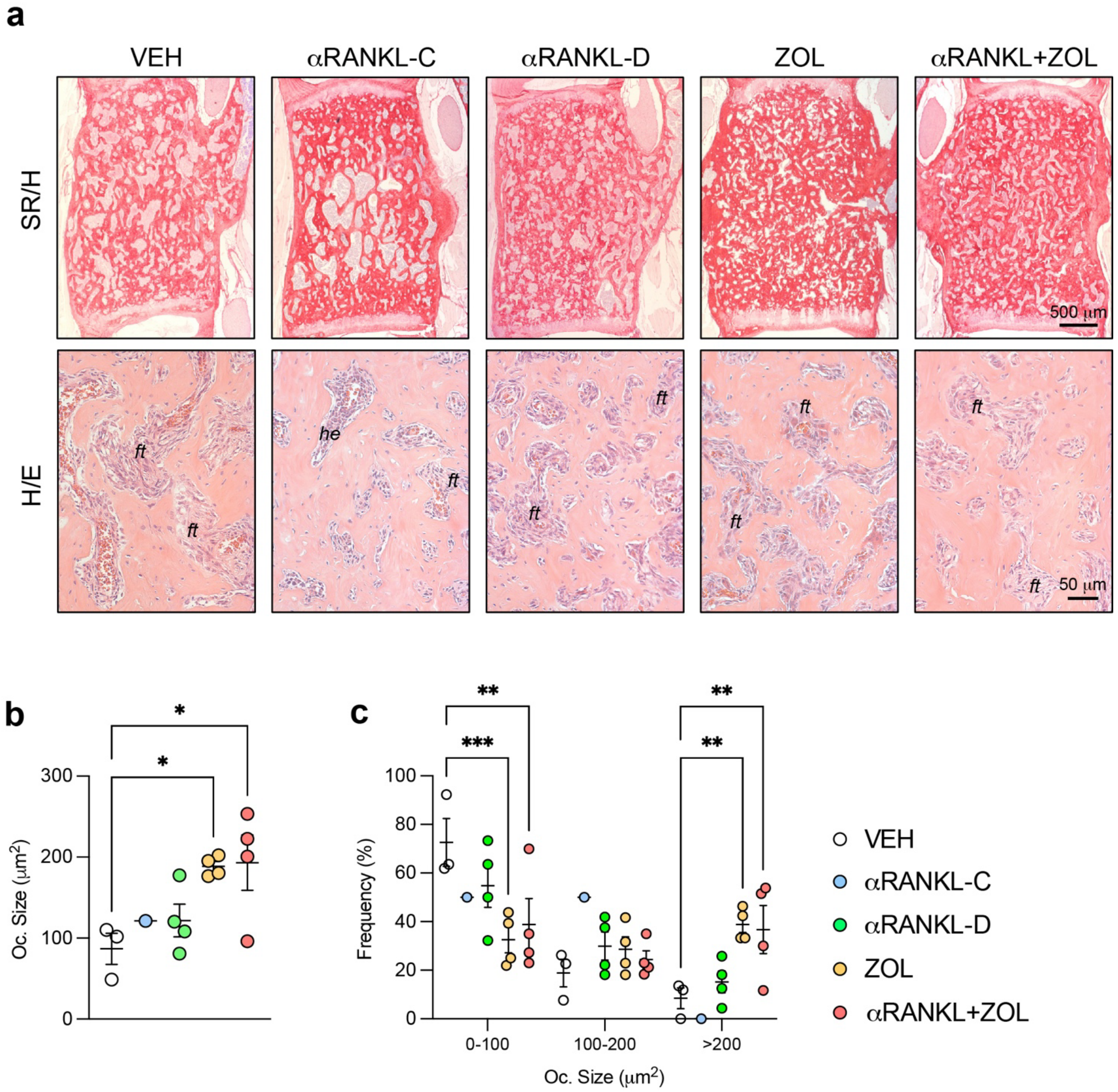


**Figure S1. Histological and histomorphometric analyses of lumbar vertebrae in treated EF1α-Gsα$^{R201C}$ mice.** a) Representative SR/H- and H/E-stained sections of lumbar vertebrae with fibro-osseous lesions. *ft* = fibrous tissue; *he* = hematopoietic marrow. b) Histomorphometric evaluation of osteoclast size (Oc. Size) in female mice. Statistical analysis with one-way ANOVA and Dunnett's multiple comparison test. c) Frequency of distribution of different osteoclast size range. Two-way ANOVA and multiple comparison test. *$p<0.05$, **$p<0.01$, ***$p<0.001$.

**Figure S2**

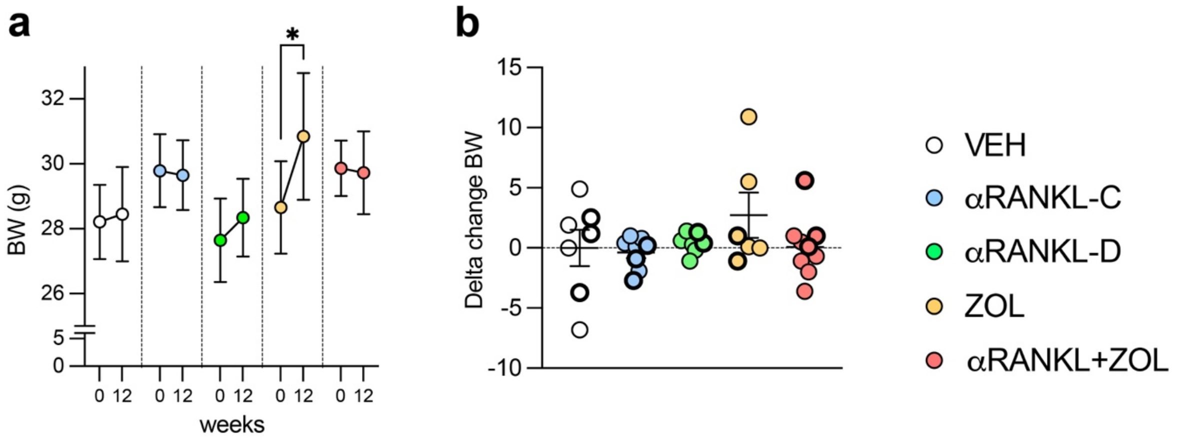


**Figure S2. Body weight changes during the treatment in EF1α-Gsα$^{R201C}$ mice.** a) Body weight values at the beginning (0) and the end (12) of treatment. Two-way ANOVA with Tukey's multiple comparison test. b) Delta change results calculated by subtracting T0 from T12 body weight.
